# Distinguishing between histories of speciation and introgression using genomic data

**DOI:** 10.1101/2022.09.07.506990

**Authors:** Mark S. Hibbins, Matthew W. Hahn

**Affiliations:** Department of Ecology and Evolutionary Biology, University of Toronto, Toronto, ON, CA M5S 3B2; Department of Biology, Indiana University, Bloomington, IN 47405; Department of Computer Science, Indiana University, Bloomington, IN 47405

## Abstract

Introgression creates complex, non-bifurcating relationships among species. At individual loci and across the genome, both introgression and incomplete lineage sorting interact to produce a wide range of different gene tree topologies. These processes can obscure the history of speciation among lineages, and, as a result, identifying the history of speciation vs. introgression remains a challenge. Here, we use theory and simulation to investigate how introgression can mislead multiple approaches to species tree inference. We find that arbitrarily low amounts of introgression can mislead both gene tree methods and parsimony methods if the rate of incomplete lineage sorting is sufficiently high. We also show that an alternative approach based on minimum gene tree node heights is inconsistent and depends on the rate of introgression across the genome. To distinguish between speciation and introgression, we apply supervised machine learning models to a set of features that can easily be obtained from phylogenomic datasets. We find that multiple of these models are highly accurate in classifying the species history in simulated datasets. We also show that, if the histories of speciation and introgression can be identified, *PhyloNet* will return highly accurate estimates of the contribution of each history to the data (i.e. edge weights). Overall, our results highlight the promise of supervised machine learning as a potentially powerful complement to phylogenetic methods in the analysis of introgression from genomic data.

## Introduction

Introgression, the process of hybridization and repeated back-crossing between previously isolated lineages, occurs frequently across the tree of life and is a common feature of modern phylogenomic datasets (Mallet et al. 2016, Taylor and Larson 2019, Dagilis et al. 2021). From a phylogenetic perspective, histories of introgression are not consistent with a strictly bifurcating phylogeny and are therefore often represented using phylogenetic networks (Huson and Bryant 2006, Huson et al. 2010, Solis-Lemus and Ane 2016, Wen et al. 2018, Wen and Nakhleh 2018). Phylogenetic networks contain additional horizontal “reticulation” edges, which are meant to display alternative histories among loci. Such networks imply that some parts of the genome follow the speciation history, while other parts follow the introgression history—a history that includes exchange between species post-speciation.

When interpreting species networks, it is often the case that one edge is assigned as the history of speciation, while the other corresponds to the history of introgression. However, methods for estimating phylogenetic networks do not explicitly model these two histories differently. For example, in Figure 1 the two networks are drawn (as is often done) to imply different species tree histories: ((A,B),C) and ((B,C),A) in Figures 1A and 1B, respectively. However, note that both networks convey the same information about a hybrid node on the lineage leading to species B, and neither contains any information about the direction of introgression. Species networks often come with estimated “inheritance” probabilities along edges connected to a hybrid node, which denote the probability that a locus in the genome follows that edge. The history of speciation between lineages is often assigned by software or by users to the edge with the higher weight (e.g. Pickrell and Pritchard 2012), under the assumption that a majority of the genome should follow this history. However, several studies have now shown that introgression can be extensive enough to affect a majority of the genome (Fontaine et al. 2015, Li et al. 2019, Forsythe et al. 2020). In such cases, phylogenetic network methods—in addition to most methods for species tree inference—will display the introgression history as the vertical history of speciation.

**Figure 1:**
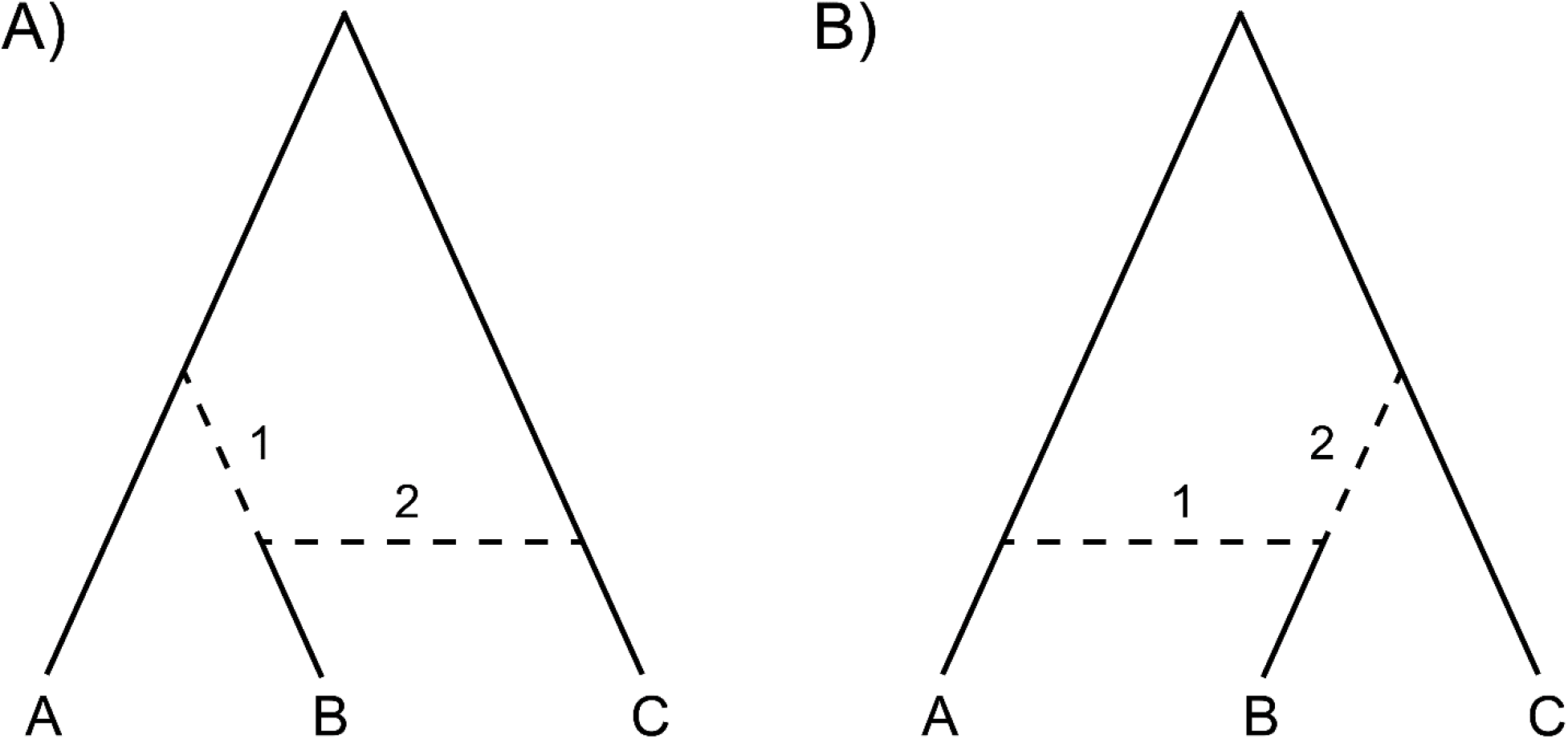
Phylogenetic networks do not explicitly distinguish histories of speciation and introgression. The networks depicted in panels A and B are drawn to imply speciation histories of ((A,B),C) and ((B,C),A) respectively, but in reality contain the same information. Both networks have a hybrid node leading to species B, with edge 1 corresponding to the history ((A,B),C) and edge 2 corresponding to the history ((B,C),A). Both networks could be estimated in a scenario where the speciation history is ((A,B),C) with post-speciation introgression between B and C, or a scenario where the speciation history is ((B,C),A) with post-speciation introgression between A and B.

Efforts to distinguish histories of speciation and introgression are complicated by incomplete lineage sorting (ILS), a stochastic process in which lineages fail to coalesce in their most recent common ancestor (Hudson 1983, Tajima 1983, Pamilo and Nei 1988). The stochastic nature of ILS means that both speciation and introgression histories can generate similar gene tree topologies at a locus. This makes it challenging to assign histories to individual loci because, for instance, the speciation history ((A,B),C) can generate a gene tree with the topology ((B,C),A) in the absence of introgression between B and C. In addition, the interaction of ILS with surprisingly little amounts of introgression can result in gene trees that match the history of introgression becoming the most common tree, misleading many standard methods for species tree inference (Eckert and Carstens 2008, Leache et al. 2014, Solis-Lemus et al. 2016, Long and Kubatko 2018, Pang and Zhang 2022). Because gene tree frequencies do not provide enough information on their own to assign one history as the speciation history, one proposal for cases with massive introgression has been to identify the introgression history as the one with the minimum average gene tree node height (Fontaine et al. 2015, Li et al. 2019, Forsythe et al. 2020). The logic behind this test is that post-speciation introgression is by definition more recent than speciation, leading to more recent coalescence. This rule should be reliable in cases with large amounts of introgression, but if introgression occurs at lower rates, then most gene trees matching the introgression history will be generated by ILS. In this case such trees will contain deeper nodes than the history of speciation. It is therefore unclear how much introgression is necessary for this approach to be useful.

While introgression and ILS reveal the complicated non-bifurcating ways that species are related to one another, understanding the history of speciation and introgression is still crucially important. The species tree provides a summary of the natural history of organisms, including their taxonomic groupings, divergence times, and dynamics of speciation and extinction (O’Meara 2012). Understanding the species history is also central to understanding the role for introgression in evolution (Rhymer and Simberloff 1996, Dowling and Secor 1997, Harrison and Larson 2014, Taylor and Larson 2019). Misspecification of the species tree can significantly impact downstream analyses, including inferences of introgression and the reconstruction of ancestral states (or the origin of novel alleles). For these reasons, distinguishing histories of speciation from those of introgression remains an important and unsolved problem.

Here, we use theory, simulation, and supervised machine learning analyses to investigate how histories of speciation and introgression might be disentangled. We ask how much introgression is necessary to mislead several approaches to species tree inference, including summary gene tree approaches, parsimony, phylogenetic network approaches, and approaches based on minimum node depths. To try to disentangle these histories, we train supervised machine learning models on simulated datasets from competing species tree topologies and introgression events, finding that the species tree can be recovered with a high degree of accuracy. Using feature importance analyses, we find that the variance in coalescence times within the two competing gene tree topologies are the most informative features, but no single piece of information alone can accurately recover the species tree topology in all areas of parameter space. Based on our findings, we provide recommendations to researchers dealing with complex histories of reticulation in phylogenomic datasets.

## Materials and Methods

### Modelling the minimum amount of introgression needed to make gene trees and site patterns supporting the history of introgression the most frequent

We begin with an exploration of a theoretical question: how much introgression is necessary to make the gene tree matching the history of introgression the most common gene tree? This question has implications for summary approaches that infer species trees using the most common gene trees among quartets or rooted triplets (e.g. Liu et al. 2010, Zhang et al. 2018), and has been addressed in different ways in a handful of previous studies. We make use of the multispecies network coalescent framework, which models the effects of introgression and ILS on gene trees simultaneously by breaking histories of speciation and introgression into separate “parent tree” histories (Meng and Kubatko 2009, Yu et al. 2014, Degnan 2018, Hibbins and Hahn 2019).

We model a rooted three-taxon tree with the species topology ((A,B),C) (Figure 2). The internal branch shared by species A and B has a length of *τ_s_,* in units of 2*N* generations. Post-speciation introgression occurs between species B and C at a rate of *δ*, which corresponds to the proportion of the genome that has introgressed. The internal branch shared by species B and C in the introgression history has a length of *τ_m_*. The value of *τ_m_* is determined by the direction of introgression (i.e. C into B or B into C) and the timing of introgression relative to speciation of A and B. Within a particular history, concordant trees (with respect to that history) arise with probability 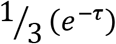, and a given discordant gene tree will arise with probability 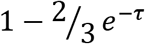, where *τ* is the length of that history’s internal branch. To get the overall expected frequencies of each gene tree topology, we weight the expected frequencies within each history by the probability that a locus follows that history (1 – *δ* for the species history, and δ for the introgression history). In what follows, we will refer to gene trees that match the species history as “AB” trees, and those that match the history of introgression as “BC” trees. For gene trees matching the species history (black tree in Figure 2B), their frequency, *f_AB_*, is

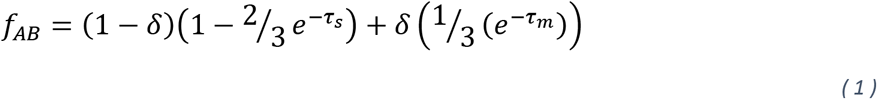

and for gene trees matching the introgression history (blue tree in Figure 2B), their frequency, *f_BC_*, is

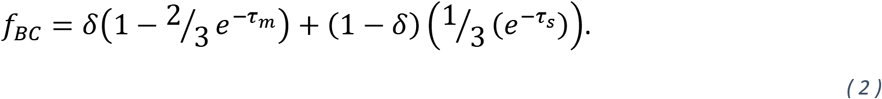

**Figure 2:**
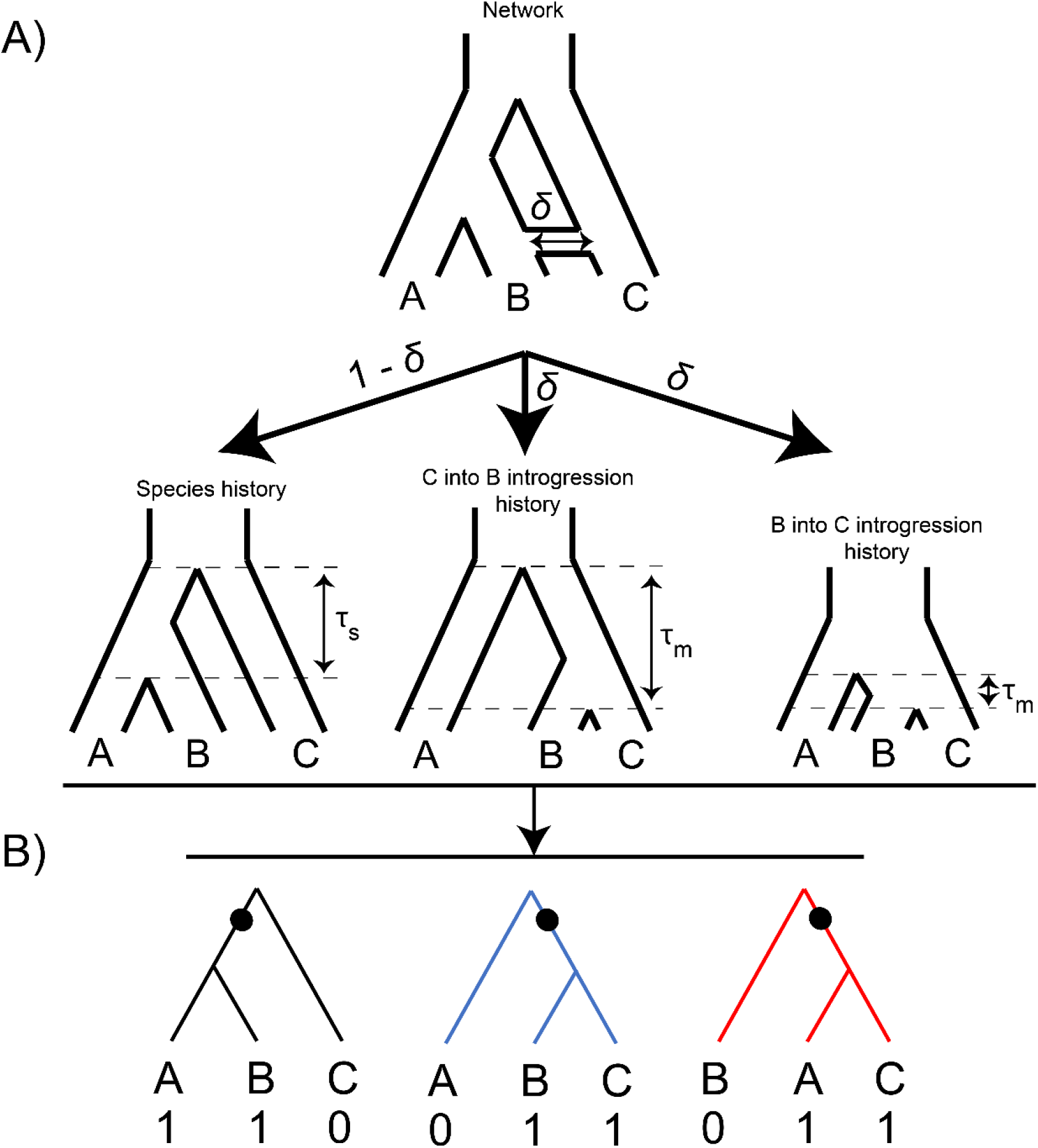
Modelling histories of speciation and introgression. A) In the multispecies network coalescent framework, a phylogenetic network can be divided into a set of possible “parent trees” which describe the histories of speciation or introgression at individual loci. Loci follow histories of introgression with probability δ, corresponding to the rate of introgression across the genome. The internal branch lengths *τ_s_* and *τ_m_*, for histories of speciation and introgression respectively, are determined by the timing of introgression relative to speciation and the direction of introgression. B) All possible histories generate gene trees under the standard multispecies coalescent model, which includes incomplete lineage sorting within each history. Mutations on the internal branches of these gene trees lead to parsimony-informative biallelic sites (shown as black dots and 0/1 ancestral/derived states at the tips).

In order to find the amount of introgression required to make *f_BC_* equal to *f_AB_*, we set the expected frequencies of the two topologies equal to one another, giving the following:

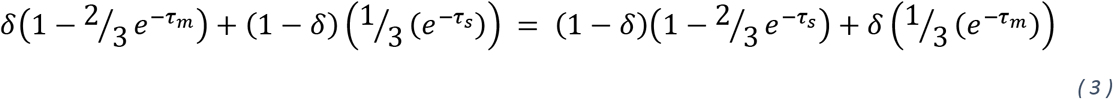

Solving for δ, we obtain the following expression:

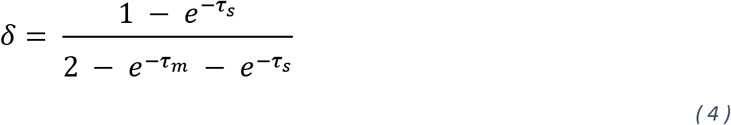

This same expression has been derived in Jiao et al. (2020) and Pang and Zhang (2022).

We used the same theoretical framework to ask a different but related question: how much introgression is necessary to make the biallelic site pattern supporting the introgression history the most common? These parsimony-informative sites are important to the performance of many standard methods for phylogenetic inference. Biallelic sites arise from mutations on internal branches of gene trees (Figure 2B). Assuming an infinite-sites model (i.e. no multiple hits), the sum of the lengths of internal branches of relevant gene trees gives us the expected number of parsimony-informative sites supporting either the ((A,B),C) or ((B,C),A) gene trees (Mendes and Hahn 2018). For gene trees that arise from lineage sorting in each history (which occurs with probability *τ* – *e*^-*τ_s_*^), the internal branch length is 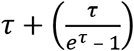 (Mendes and Hahn 2018), while for all other gene trees (i.e. the ones due to incomplete lineage sorting in any history) the internal branch length is 1 (in units of *2N* generations). To obtain the average internal branch length across histories, we weight the relevant branch lengths within each history by the frequencies of the gene trees, and then across histories by their probabilities at a locus. This gives the following expression for biallelic sites that support the species history:

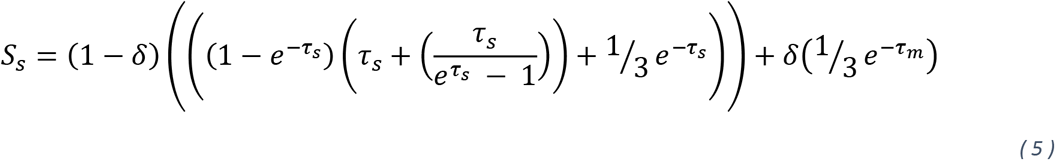

and this expression for sites supporting the history of introgression:

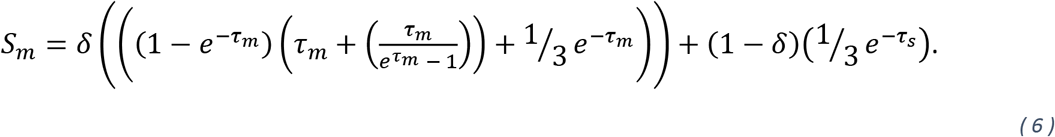

To find the *δ* value required to make site patterns from both histories equally frequent, we set these two expressions equal, as before. After solving for *δ* and simplifying, we obtain the following expression:

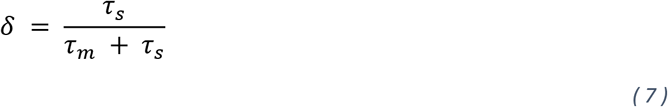

To our knowledge, this relationship has not been found before.

### Modelling the minimum gene tree node height

We also used our model to investigate the “minimum node height” criterion for distinguishing between histories (Fontaine et al. 2015, Li et al. 2019, Forsythe et al. 2020). For this analysis we make use of the expected coalescence times and conditional gene tree frequencies derived in Hibbins and Hahn (2019). Since gene trees in real data arise from a combination of different histories, we obtain general expressions here by averaging the time to first coalescence across all possible gene trees that share the same topology. These times to first coalescence are measured conditional on a gene tree topology, so they must be weighted by the frequency of a particular gene tree relative to the frequency of all other possible gene trees that share that topology, rather than simply the overall expected frequency. We used these expressions to calculate the expected node heights of AB and BC gene trees over a range of parameter values from our model, and asked in which parts of the parameter space each node height was the smallest. See Section 1 of the Supplementary Materials and Methods for the relevant expressions.

### *Simulating gene trees with* msprime

To validate our theoretical results, and to conduct downstream analyses, we simulated gene trees under various speciation and introgression scenarios using the Python package *msprime* (Baumdicker et al. 2022). For each simulated dataset, we denoted the history of speciation among lineages as a Newick string and then used *Demography.from_species_tree()* to convert this into an *msprime* demography object. Introgression was added to these demography objects using the *demo.add_mass_migration* function with the specified timing, direction, and rate of introgression. We then simulated tree sequences using *sim_ancestry*(), specifying a single haploid sample from each lineage with a recombination rate of 1×10^-8^ and a sequence length of 1×10^7^. These tree sequences were then converted into Newick trees for further parsing. We converted branch lengths to units of 2*N* generations assuming *N* = 10000, and sampled every third gene tree to reduce the effects of spatial autocorrelation, resulting in approximately 1200 gene trees per dataset (i.e. for each unique combination of parameters).

### Supervised machine learning analyses to predict the history of speciation

To investigate whether it is possible to accurately distinguish histories of speciation and introgression, we applied supervised machine learning approaches to simulated datasets using the Python package *scikit-learn* (Pedregosa et al. 2011). Using the approach described in the previous section, we simulated gene tree datasets in *msprime* under two competing histories: one where A and B are most closely related with post-speciation introgression between B and C, and one where B and C are most closely related with post-speciation introgression between A and B. The histories of speciation in these two scenarios were used as labels for binary classification. We simulated a 10×10×10 grid of *τ_s_, τ_m_,* and *δ* values, in each direction of introgression and for each possible history, resulting in 4000 total simulated datasets. For each history, the two simulated directions of introgression were “P3 into P2” for introgression from the unpaired lineage into the paired lineage, and “P2 into P3” for introgression from the paired lineage into the unpaired lineage. These two directions correspond to C into B and B into C introgression respectively for the history of BC introgression, and A into B and B into A introgression respectively for the history of AB introgression. For each of these, we used *ete3* (Huerta-Cepas et al. 2016) to parse 21 features, including the frequency of each of the three gene trees, the time to coalescence of each pair of species within each of the gene trees, and the variances of these times to coalescence. We split simulations into a training dataset of 3000 observations and a test dataset of 1000 observations, and then standardized all features using *scikit-learn*’s *StandardScaler()*. We trained five binary classification models on the training dataset, all using the default model parameters: logistic regression, support vector machine, Gaussian naïve Bayes, decision tree, and random forest. Finally, the performance of each of these trained models was scored on the test set.

We assessed the importance of each feature to each model’s predictive power on the test set using the *permutation_importance()* function in *scikit-learn*. This analysis randomly permutes each feature in the dataset and scores its importance by how much this permutation decreases model performance. Permutation was repeated 30 times for each feature, and a feature was deemed significantly important for a model if its mean importance score across replicates was at least twice that score’s standard deviation. We ranked the importance of each feature across models by taking the average importance score.

### *Assessment of* PhyloNet *edge weight estimates*

We ran *PhyloNet*’s *InferNetwork_ML* method (Yu et al. 2014) on simulated gene tree datasets to evaluate the accuracy of edge weight estimates against simulated introgression proportions. We simulated gene trees with *msprime* under the same parameter combinations as for the machine learning analyses, and used a custom python script to convert the outputs of these simulations into NEXUS-formatted input files for *PhyloNet.* On the *PhyloNet* outputs, we used the Julia package *PhyloNetworks* (Solis-Lemus et al. 2017) and Python package *ete3* (Huerta-Cepas et al. 2016) to extract the weight of the edge in the network that corresponds to the known history of introgression. In cases where no introgression was inferred by *PhyloNet*, we assigned a weight of 0 when the estimated history corresponded to the species tree, and a weight of 1 when itcorresponded to the introgression history. Edge-weight estimates for each simulated value of the rate of introgression were averaged across all combinations of branch length parameters.

### Data Availability

Scripts and data for all analyses are available at https://github.com/mhibbins/dist_histories.

## Results

### Arbitrarily small amounts of introgression can make gene trees and site patterns supporting the history of introgression the most frequent

To visualize the amount of introgression necessary to make discordant gene trees more common than concordant ones, we plotted the *δ* value derived in Equation 4 over the space of *τ_s_* and *τ_m_* (Figure 3A, B). Overall, we find that the smaller *τ_s_* is, the less introgression is necessary to make the introgression tree the most common gene tree. This occurs because smaller values of *τ_s_* lead to more ILS (and therefore more discordance), even in the absence of introgression. As the amount of discordance approaches 66% in the species tree history (i.e. a star tree), the amount of introgression required to tip the balance toward the topology matching the introgression history approaches 0. The minimum value of *δ* is also affected by the timing and direction of introgression, with more recent introgression from C into B resulting in a lower minimum amount required. Both the timing and direction of introgression affect the amount of ILS within introgressed histories, as more recent introgression (i.e. larger τ_m_) and introgression from C into B both lead to longer internal branches in the introgressed history (Figure 2). Conversely, if ILS is low in the species history (large *τ_s_*) and high in the introgression history (small *τ_m_* and B into C introgression), rates of introgression can be very high (approaching 100%) and still not result in topologies matching the introgression history being most common. The histories of speciation and introgression in the extremes of the parameter space considered here are summarized in Figure 3D.

**Figure 3:**
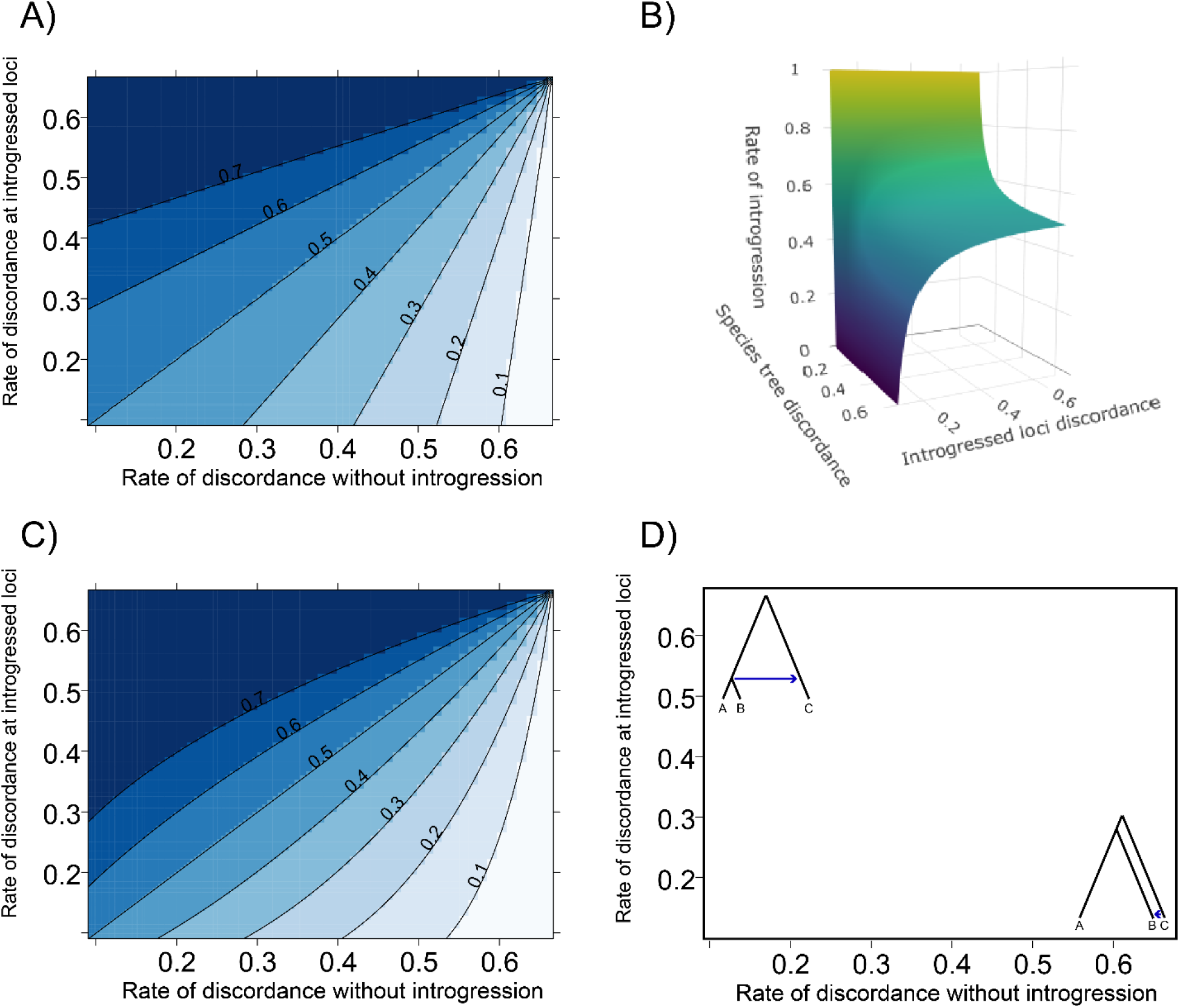
The amount of the genome affected by introgression that is necessary to make gene trees (panels A and B) and site patterns (panel C) matching the introgression history the most common, as a function of the rate of discordance with respect to both histories. In panels A and C, the contours show the minimum necessary rate of introgression in that part of parameter space. Panel B plots the result in panel A as a surface with the rate of introgression as the vertical axis. Panel D summarizes the continuum of possible speciation and introgression histories across the parameter space.

When plotting the minimum *δ* value for parsimony-informative sites derived in Equation 7, we find a similar overall pattern to the result for gene tree frequencies, but with a slightly different shape to the contours (Figure 3C). Once again, large amounts of ILS in the species history mean that little introgression is necessary to make sites supporting the introgression history the most common. The differences between Figures 3A and 3C imply that parsimony methods may perform better than summary methods under certain speciation and introgression scenarios, and vice versa. To better visualize these differences, we plot the gene tree frequency-based *δ* value subtracted from the parsimony-based *δ* value in Supplementary Figure 1. In general, we find that methods that use gene tree frequencies should perform better (i.e. require more introgression to be misled) than parsimony methods when the *δ* value required is less than 50%, while parsimony methods should perform better when the *δ* value is greater than 50%. The areas where the two methods have the greatest difference in performance are not linear with the amount of discordance, but rather fall in regions with intermediate levels of discordance. In these parts of parameter space, the difference in minimum *δ* between methods is as high as approximately 11%. This means, for example, that when *δ* is less than 50%, summary methods can tolerate an 11% higher rate of introgression than parsimony methods before the introgression tree becomes most common. In either case, little introgression is needed to “hide” the species tree when there is a lot of ILS in the species history.

The results presented in this section suggest that it may be difficult to distinguish between histories using gene tree frequencies or informative sites alone, even under relatively modest introgression scenarios. Previous studies have proposed using the minimum average node height among gene tree topologies to distinguish the species history from the introgression history (Fontaine et al. 2015, Li et al. 2019, Forsythe et al. 2020). Using the same modelling framework as the previous section, we evaluated the space over which the “minimum node height” criterion may be effective at inferring the introgression history. We found that the minimum amount of introgression required to make the minimum average node height correspond to the history of introgression, rather than that of speciation, varied from 20% to approximately 40% (Figure 4). This variation depends primarily on the timing of introgression relative to speciation, with more recent introgression resulting in less introgression required. The direction of introgression has little effect; Figure 4 shows the result for equal amounts of introgression in both directions, while Supplementary Figure 2 shows the pattern for each direction separately. As the species history approaches a star phylogeny, the necessary rate can fall below 20% and approaches 0 (but this is a relatively small part of the parameter space). These results suggest that the average minimum node height can be informative when the rate of introgression is high, but will fail in large parts of parameter space. Overall, it seems unlikely that any one feature alone will reliably help to identify one history over another.

**Figure 4:**
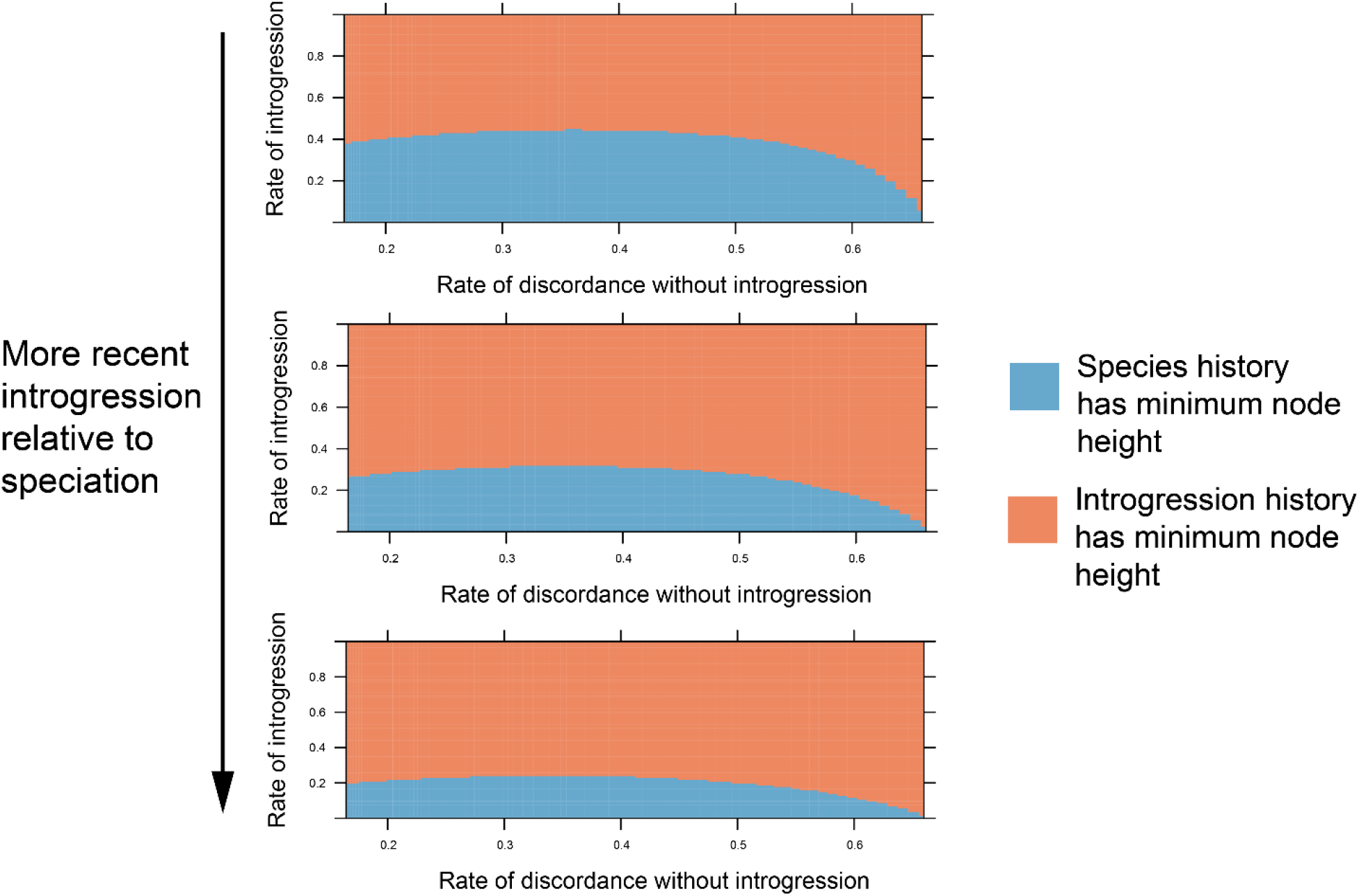
Distinguishing histories of speciation and introgression using the minimum gene tree node height. In blue areas, the minimum node height (averaged across all gene trees sharing a topology) is in gene trees matching the species history, while in orange it is in gene trees matching the introgression history. The boundary between the two denotes the minimum amount of introgression necessary to make the introgression history the one with the shortest node height. This minimum amount of introgression decreases as the species tree approaches a star tree (i.e. as the rate of discordance without introgression increases) and as introgression becomes more recent relative to speciation (from top to bottom, *t*_m_ values of 0.45, 0.3, 0.15, respectively). In most of parameter space, a significant amount of introgression is necessary, ranging from 20% (bottom) to 40% (top). Patterns shown are for equal rates of introgression in both directions.

### Supervised machine learning recovers the history of speciation with high accuracy

Supervised machine learning is a powerful tool for data classification problems, and its applications in population genetics and phylogenetics are growing (Schrider and Kern 2018). Supervised machine learning methods can efficiently use multidimensional inputs in order to make accurate classification of a dataset. Here, we made use of supervised machine learning for binary classification of the history of speciation in simulated datasets. We used *msprime* (Baumdicker et al. 2022) to simulate datasets containing two competing histories: one where the species tree is ((A,B),C) with introgression between C and B, and one where the species tree is ((B,C),A) with introgression between A and B (Figure 1). Simulations were conducted for parameter combinations over the space of our model for both possible histories. These analyses consider the two possible directions of introgression separately, which we refer to hereafter as “P3 into P2” for C into B or A into B introgression, and “P2 into P3” for B into C or B into A introgression. From each simulated dataset we extracted 21 features, including gene tree frequencies, average coalescence times, and variances in coalescence times within each topology. These features were passed to five machine learning models for binary classification implemented in *scikit-learn* (Pedregosa et al. 2011). After training, we used 1000 test datasets to see if the models could accurately classify a dataset as coming from one of the two competing species histories. We found that the supervised machine learning methods recovered the history of speciation with high accuracy, ranging from 76.3% for naïve Bayes to 93.3% for a random forest classifier (Table XX).

**Table 1:** Performance of our supervised machine learning classifiers on simulated test datasets.

| **Model** | **Performance score on test set** |
| --- | --- |
| Logistic regression | 0.792 |
| Support vector machine | 0.913 |
| Gaussian Naïve Bayes | 0.763 |
| Decision tree | 0.878 |
| Random forest | 0.933 |

To understand which features were the most predictive of the correct species history across models, we conducted a permutation feature importance analysis as implemented in *scikit-learn*. This analysis randomizes individual features in the dataset one at a time and evaluates how this impacts the model’s predictive accuracy. We ranked each feature by its overall importance score summed across the five models. We found that most features (17/21) had significant importance scores in at least one model, and there was notable variation in the importance of features across models (Figure 5). Across models, the most predictive features were the variances in coalescence times of the sister species in gene trees matching the two relevant histories: i.e. the variance in time to coalescence of A and B in AB gene trees (AB_AB_var in Figure 5) and B and C in BC gene trees (BC_BC_var). The next four most important features (in most models) were the node distances in those gene trees (AB_AB_dist and BC_BC_dist) and the frequencies of the gene trees (AB_freq and BC_freq). The importance of other features after these six is notably lower and varies between models.

**Figure 5:**
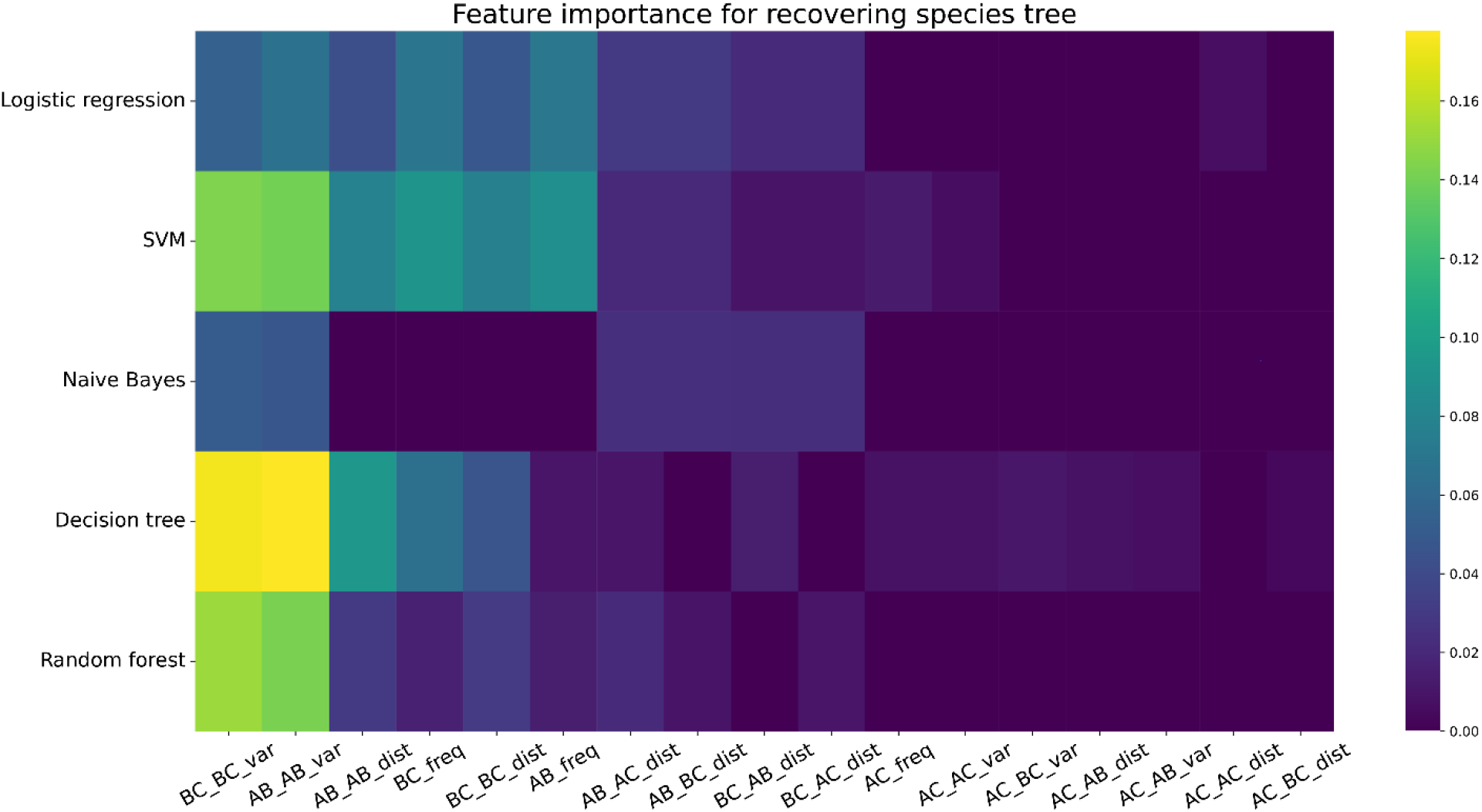
Model features (x-axis) ranked by importance score (color scale) across all our trained classification models (y-axis). In feature names, the first pair of letters indicate the gene tree topology, and the second pair indicate the pair of species within that gene tree topology, so the feature “AB_AC_dist” is the branch length distance between species A and C in gene trees where A and B are sister taxa. Four features are not shown which did not have a significant importance score in any of our trained models.

To help uncover general patterns that might be useful for distinguishing between competing histories in real datasets, we plotted the behavior of the six most important features discussed in the previous paragraph in each of the two histories in our simulated datasets (Figure 6). Each variable is plotted over the space of *δ* values for each direction of introgression and averaged across all values of *τ_s_* and *τ_m_* within a value of *δ.* Generally, the gene tree with the lower variance in the time to first coalescence corresponds to the species history. This pattern remains true except when rates of introgression are very low (less than approximately 5%) or very high (greater than approximately 75%). The consistency of this pattern across a wide range of possible *δ* values helps to explain why these features emerged as the most important in our machine learning models (Figure 5). The other features behave as expected from our theoretical results, with increasing rates of introgression decreasing the minimum node height and increasing the frequency of trees that match the introgression history. In general, the minimum node height corresponds to the introgression history when the rate of introgression is higher than ~25%, and the most common gene tree matches the species history when the rate of introgression is less than 50%, but there is significant deviation from these patterns depending on rates of discordance within each history, as well as the direction of introgression (Supplementary Figure 3). Nonetheless, the large differences in values between histories in most of the space makes these features highly informative together or when combined with other information.

**Figure 6:**
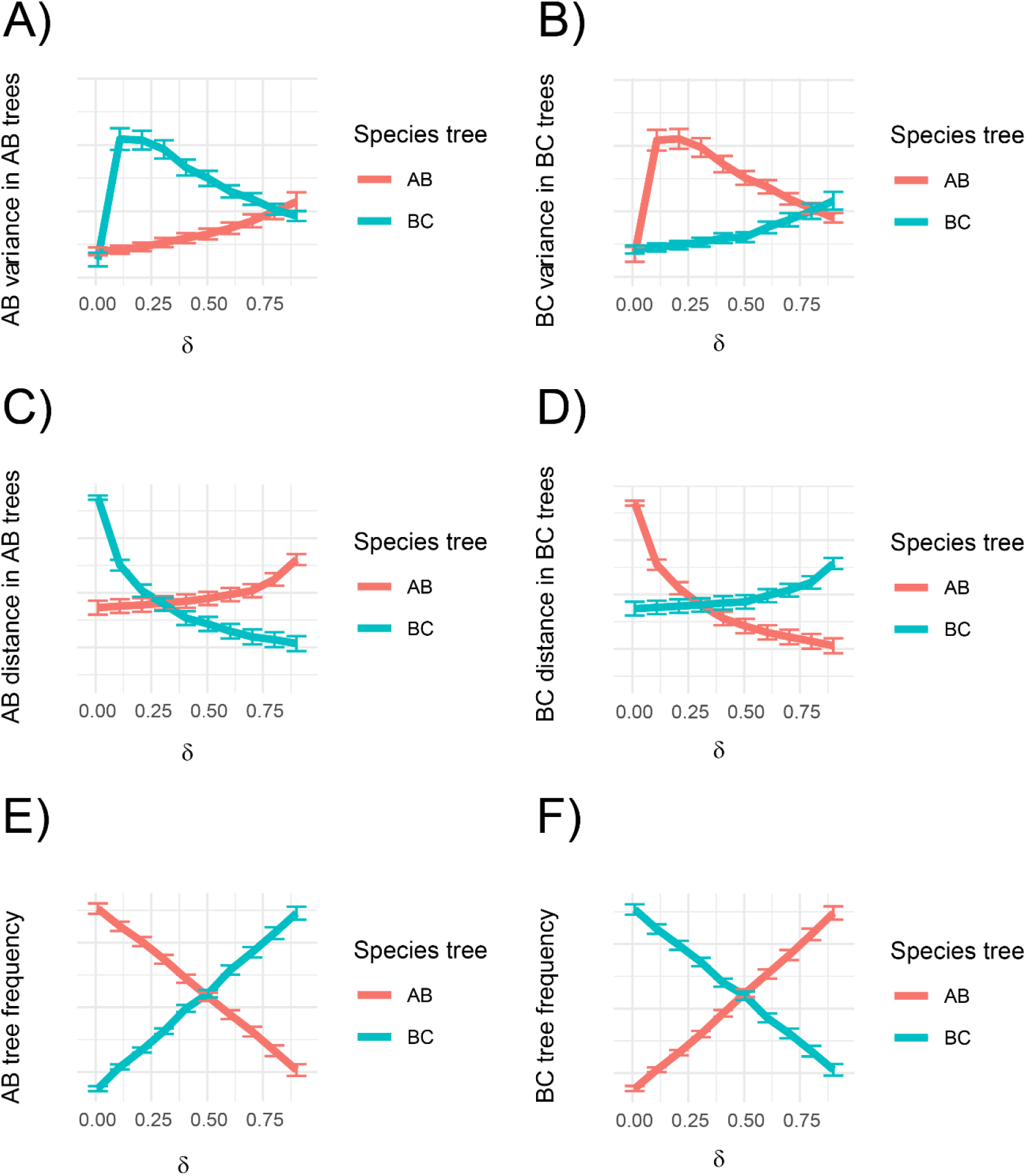
Behavior of the six most informative features identified from our machine learning classifiers, for P3 into P2 introgression (C into B or A into B, depending on which history is the speciation history and which the introgression history). Color denotes the true history of speciation underlying the simulated data, and the error bars denote the standard error introduced by variation in *τ_s_* and *τ_m_* within each value of *δ* on the x-axis. A) The variance in the time to coalescence of A and B in AB gene trees (denoted *AB_AB_var* in Figure 5). B) The variance in the time to coalescence of B and C in BC gene trees (denoted *BC_BC_var* in Figure 5). C) The average time to coalescence of A and B in AB gene trees (denoted *AB_AB_dist* in Figure 5). D) The average time to coalescence of B and C in BC gene trees *(BC_BC_dist* in Figure 5). E) The frequency of AB gene trees (denoted *AB_freq* in Figure 5). F) The frequency of BC gene trees (denoted *BC_freq* in Figure 5).

### PhyloNet accurately estimates edge weights for different histories in the presence of misleading gene tree frequencies

While the machine learning results in the previous section provide possibilities for distinguishing histories of speciation and introgression, they do not provide information on how much of the genome corresponds to each history. Phylogenetic network methods do not explicitly label histories, but they may nonetheless be able to accurately estimate the edge weights corresponding to these histories (which should correspond to *δ*). These estimates may be especially useful in parts of the parameter space where gene tree frequencies do not correspond intuitively to introgression probabilities. For example, in the top left space of Figure 3A and C, introgression probabilities can be as high as 70% or more, though the gene tree matching the species history is still the most common. To assess edge weights, we simulated gene tree datasets with *msprime* and then passed them to the *InferNetwork_ML* method of *PhyloNet* (Yu et al. 2014) and asked whether the estimated reticulation edge weights reflected gene tree frequencies or the true simulated rates of introgression.

In general, we found that *PhyloNet*’s estimated edge weights correspond to the actual proportions of speciation and introgression histories, rather than to gene tree frequencies (Figure 7). We observed substantial variation in the frequency of BC gene trees, especially at high rates of introgression, where essentially any frequency is possible depending on the values of other parameters. This is especially true when introgression was from B into C: for example, with a rate of introgression of 90%, the average frequency of BC gene trees was only ~45%, with a large standard error (Figure 7, bottom right). This is in line with the theoretical results plotted in Figure 3A and 3B. Despite this, *PhyloNet* is generally able to accurately estimate edge weights, with a nearly 1:1 correspondence between the simulated amount of introgression and the mean estimated amount (though there is also significant variation in these estimates). Overall, these results suggest that if the histories of speciation and introgression can be correctly identified, *PhyloNet* is able to accurately estimate what proportion of the genome corresponds to each history.

**Figure 7:**
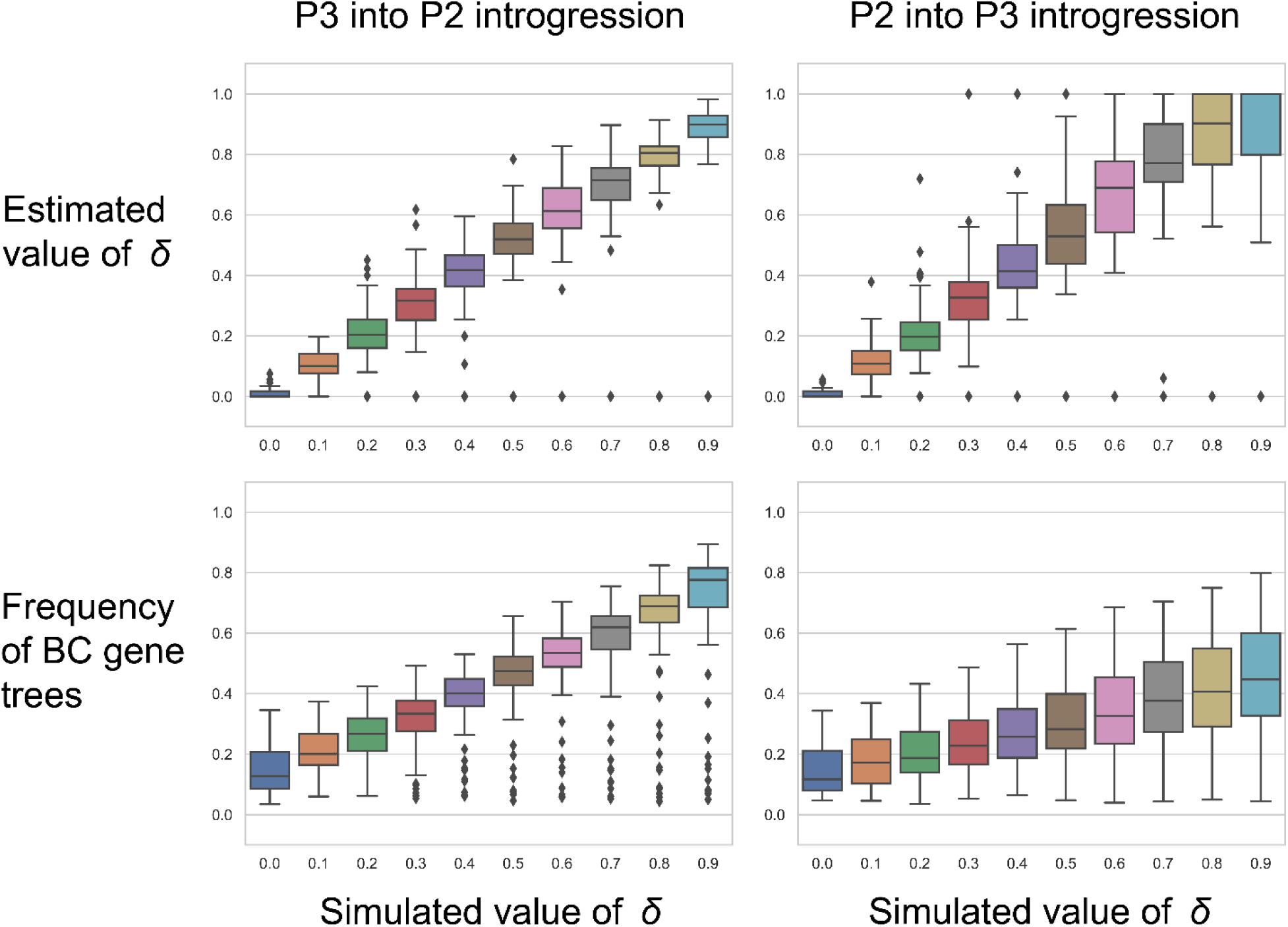
Ability of *PhyloNet* to estimate the proportion of the genome that has introgressed (top row) relative to observed gene tree frequencies (bottom row). For each value of *δ,* estimates are grouped across all simulated values of *τ_s_* and *τ_m_.* Individual points show outlier estimates; estimates of 0 are assigned when the species tree is inferred with no reticulations, and estimates of 1 are assigned when the introgression history is inferred with no reticulations.

## Discussion

Genomes often contain a complex mosaic of different relationships among the same set of species. Understanding the historical processes that generate these different histories, such as speciation and post-speciation introgression, is critical to obtaining a complete understanding of the natural history of organisms. Here, we have used theory, simulation, and a supervised machine learning approach to show that while currently available approaches to species tree inference may often be misled in the presence of introgression, it should be possible in principle to distinguish among these histories using the information contained in genome-scale datasets. By exploring the behavior of the most informative features in our machine learning models, in addition to highlighting the accuracy of edge-weight estimates in phylogenetic network approaches, we can now provide recommendations for future analyses and methods, as well as discussing the assumptions and limitations of our work.

For the sake of tractability, our theoretical model (and simulation analyses) makes simplifying assumptions that have implications for patterns in genomic data. First, we assume a single, discrete, post-speciation introgression event between one pair of non-sister taxa. There are a multitude of ways that real introgression scenarios could be more complicated, including multiple discrete events between different pairs of species, or continuous periods of gene flow rather than discrete pulses of hybridization. As long as introgression is primarily between one pair of lineages, the general patterns of genomic features we have reported here should hold, but the power of particular features to distinguish between different histories may be affected. Alternatively, if other lineages are involved in introgression scenarios, this may affect gene tree patterns in ways that are not accounted for by our model. For example, introgression between ancestral sister lineages can result in *both* discordant gene tree topologies becoming more common than the species tree topology, rather than just one (Solis-Lemus et al. 2016, Long and Kubatko 2018, Jiao and Yang 2021). Additionally, introgression from a more distantly related ghost lineage can cause an incorrect history of introgression to be inferred from available data (Tricou et al. 2022), resulting in loci matching the “introgression history” having longer branches than expected. Lastly, introgression events between multiple pairs of lineages will present a more complex problem than simply distinguishing between two competing histories, as new histories are introduced by each event; future work will be needed to address such scenarios.

Our theoretical results suggest that standard methods for species tree inference will often fail to infer the correct species tree in the presence of introgression. If levels of ILS in the species history are sufficiently high, very little introgression is necessary to mislead both summary and parsimony methods into inferring the introgression history as the species tree. This is important because both approaches have been shown to outperform maximum-likelihood inference in the presence of discordance due to ILS (Mirarab et al. 2016, Mendes and Hahn 2018) and are often applied to datasets with high rates of gene tree discordance. We predict that maximum likelihood methods would be similarly misled. While phylogenetic network approaches consider both ILS and introgression as sources of discordance, they do not explicitly label these histories. In our simulated datasets, *PhyloNet* frequently inferred the introgression history as the only history with no reticulations when the simulated rate of introgression was greater than 70%. Overall, these findings highlight the need for methods of species tree inference than can better accommodate introgression.

In addition to its implications for species tree inference, our model provides a cautionary note on the utility of gene tree frequencies in other introgression-related inferences. The frequency of gene trees matching the introgression history does not always correspond neatly to the rate of introgression across the genome, owing to ILS at introgressed loci. This is highlighted by the fact that the *D*-statistic (Green et al. 2010), a test based on gene tree frequencies, is a poor estimator of the rate of introgression (Martin et al. 2015, Hamlin et al. 2020). In addition, the lack of information in gene tree frequencies complicates inferences of homoploid hybrid speciation, a controversial process (Schumer et al. 2014, Nieto Feliner et al. 2017, Schumer et al. 2018) in which hybridization is proposed to cause speciation without a change in ploidy. One potential piece of evidence for homoploid hybrid speciation is equal frequencies of the two most common gene tree topologies across the genome, which one might expect if reproductive isolation begins immediately in F1 hybrids. Our results show that it is possible to have two equally frequent gene trees at any rate of introgression (Figure 3), not only with *δ* =0.50. This highlights the difficulty in devising tests for hybrid speciation based on only gene tree frequencies. Fortunately, we found that *PhyloNet* accurately estimates the rate of introgression across parameter space of our model, demonstrating the utility of additional branch-length information in estimating the amount of introgression.

Our machine learning models were able to classify the species history of simulated datasets with high accuracy, especially models based on decision trees. This suggests that it may be possible to build a similar classifier that can be applied to real datasets in cases where there are two competing histories in the data. One hurdle to developing such an approach, however, is the necessity of generating simulated training datasets. These simulations require knowledge of demographic parameters that cannot necessarily be accurately estimated if the histories of speciation and introgression are not known. One solution is to simulate over a large space of possible parameter combinations that covers the entire range of biologically plausible values, and then to build a classifier from this data for application to empirical data. Such a classifier could easily be applied to larger phylogenies if there is only a single introgression event; for multiple intogression events, it may be necessary to divide larger trees into a set of classification tasks for three-taxon subtrees. Similar simulation approaches to training machine learning classifiers have found success in application to real datasets for other introgression-related questions. For example, Schrider et al. (2018) identified introgressed loci in two *Drosophila* species by simulating genomic windows and training a classifier on summary statistics. Burbrink and Gehara (2018) identified ancient introgression in the phylogeny of New World kingsnakes using a classifier trained on pairwise distance features obtained from simulated gene trees. One other possibility is to use unsupervised machine learning approaches, such as a generative adversarial network (e.g. Wang et al. 2021), to generate histories of speciation and introgression with gene tree features that most closely resemble those observed in the empirical data.

Our feature importance analysis revealed general patterns in genomic summary statistics that may be useful for distinguishing among histories. The behavior of the six most informative features is in line with expectations from our mathematical model and other previous work, and demonstrates how these features can each reveal different pieces of information about the histories of the data. The most useful features were the variances in coalescent times within gene trees, as they are informative over a large part of the space of possible *δ* values. Gene trees matching the introgression history tended to have a higher variance in coalescence times; this is consistent with introgression introducing a more recent lower bound to coalescence, increasing the range of possible coalescence times. This lower bound also explains the informativeness of gene tree pairwise distances, which can take smaller possible values in gene trees matching the introgression history. Nevertheless, no single feature is completely informative in all areas of parameter space. This observation explains why most of our features had significant importance scores in at least one model. For example, the variables *AB_AC_dist* and *BC_AC_dist* should be informative about the direction of introgression, since P2 into P3 introgression affects the time to coalescence of A and C (Hibbins and Hahn 2019). These results highlight the power of machine learning to combine different pieces of information to make accurate predictions.

Overall, our work highlights the challenges of distinguishing between speciation and introgression histories, but also provides promising paths forward that may eventually lead to the development of methods for carrying out this task. While massive amounts of introgression may break down the distinctions between contemporary species, understanding the historical processes of speciation and hybridization underlying these lineages is still crucial for studying macroevolutionary processes such as speciation and extinction, as well as the evolution of traits over time.

## Acknowledgements

We would like to thank Claudia Solís-Lemus and Marianne Bjorner for help with *PhyloNetworks.* This work was supported by National Science Foundation grant DEB-1936187 awarded to Matthew Hahn, and the EEB Postdoctoral Fellowship awarded to Mark Hibbins by the Department of Ecology and Evolutionary Biology at the University of Toronto.

## Notes

https://github.com/mhibbins/dist_histories

### Competing Interest Statement

The authors have declared no competing interest.

