## Supplement Materials and Methods for "Distinguishing between histories of speciation and introgression using genomic data"

### Supplementary Materials and Methods

#### 1 Theory for gene tree node heights

To investigate the utility of the "minimum node height" criterion for identifying the introgression history, we modelled expected coalescence times in gene trees using the expectations derived in Hibbins and Hahn (2019). Specifically, we are interested in the expected time of the first coalescence event in AB and BC gene trees, which match the histories of speciation and introgression, respectively (see Figure 2 in main text). To obtain these expectations, we must write the expectation for each possible gene tree within a topology that can be produced across the two histories. In broad terms, the expected node heights can therefore be written as follows:

$$h_{AB}|AB = f_{AB1_1}E[t_{A-B}|AB1_1] + f_{AB2_1}E[t_{A-B}|AB2_1] + f_{AB_2}E[t_{A-B}|AB_2] + f_{AB_3}E[t_{A-B}|AB_3]$$

for the height of A and B in AB gene trees, and

$$h_{BC}|BC = f_{BC1_2}E[t_{B-C}|BC1_2] + f_{BC2_2}E[t_{B-C}|BC2_2] + f_{BC_1}E[t_{B-C}|BC_1] + f_{BC1_3}E[t_{B-C}|BC1_3] + f_{BC2_3}E[t_{B-C}|BC2_3]$$

for the height of B and C in BC gene trees. For AB gene trees, there are four trees to consider: $AB1_1$ and $AB2_1$ arise from lineage sorting and incomplete lineage sorting in the species history, respectively; $AB_2$ arises from ILS in a history of C into B introgression, and $AB_3$ from ILS in a history of B into C introgression. For BC gene trees, there are five trees: $BC1_2$ and $BC2_2$ for lineage sorting and ILS in a history of C into B introgression; $BC1_3$ and $BC2_3$ for lineage sorting and ILS in a history of B into C introgression; and $BC_1$ for ILS in the species history.

Since we are modelling coalescence times, we must use separate variables for the time of each speciation and introgression event, rather than an internal branch variable. We define $t_2$ as the time of speciation between C and the ancestor of A and B, $t_1$ as the time of speciation of A and B, and $t_m$ as the timing of the introgression event between B and C. In general, the first coalescence event between any two of three lineages in a single population is expected to occur $2/3N$ generations in the past, while coalescence of the remaining two lineages takes $2N$ generations. The time to first coalescence in lineage sorting trees has a more complex form, as it is conditional on coalescence occurring before $t_2$. With these conventions, and labelling the time to coalescence (for example) between lineages A and B in a specific topology as $t_{A-B}$, we have the following set of equations for the expected times of the first coalescent event:

$$E[t_{A-B}|AB1_1] = t_1 + \left(1 - \frac{t_2 - t_1}{e^{(t_2-t_1)} - 1}\right)$$

$$E[t_{A-B}|AB2_1] = E[t_{A-B}|AB_2] = t_2 + \frac{1}{3}$$

$$E[t_{A-B}|AB_3] = t_1 + \frac{1}{3}$$

$$E[t_{B-C}|BC1_2] = t_m + \left(1 - \frac{t_2 - t_m}{e^{(t_2-t_m)} - 1}\right)$$

$$E[t_{B-C}|BC2_2] = E[t_{B-C}|BC_1] = t_2 + \frac{1}{3}$$

$$E[t_{B-C}|BC1_3] = t_m + \left(1 - \frac{t_1 - t_m}{e^{(t_1-t_m)} - 1}\right)$$

$$E[t_{B-C}|BC2_3] = t_1 + \frac{1}{3}$$

(Hibbins and Hahn 2019).

Each of these times to coalescence are weighted by a frequency term. Since the node heights of interest are conditional on a specific gene tree topology, these frequency terms must be normalized so that they represent a proportion of all the gene trees sharing that topology, rather than a proportion of all possible gene trees. Using $\delta_2$ as the rate of C into B introgression, and $\delta_3$ as the rate of B into C introgression, this gives the following set of expressions:

$$f_{AB1_1} = \frac{(1-\delta_2 - \delta_3)\left(1 - e^{-(t_2-t_1)}\right)}{(1-\delta_2 - \delta_3)(1-e^{-(t_2-t_1)}) + (1-\delta_2 - \delta_3)\frac{1}{3}e^{-(t_2-t_1)} + \delta_2\frac{1}{3}e^{-(t_2-t_m)} + \delta_3\frac{1}{3}e^{-(t_1-t_m)}}$$

$$f_{AB2_1} = \frac{(1-\delta_2 - \delta_3)\frac{1}{3}e^{-(t_2-t_1)}}{(1-\delta_2 - \delta_3)(1-e^{-(t_2-t_1)}) + (1-\delta_2 - \delta_3)\frac{1}{3}e^{-(t_2-t_1)} + \delta_2\frac{1}{3}e^{-(t_2-t_m)} + \delta_3\frac{1}{3}e^{-(t_1-t_m)}}$$

$$f_{AB_2} = \frac{\delta_2\frac{1}{3}e^{-(t_2-t_m)}}{(1-\delta_2 - \delta_3)(1-e^{-(t_2-t_1)}) + (1-\delta_2 - \delta_3)\frac{1}{3}e^{-(t_2-t_1)} + \delta_2\frac{1}{3}e^{-(t_2-t_m)} + \delta_3\frac{1}{3}e^{-(t_1-t_m)}}$$

$f_{AB_3}$

$$= \frac{\delta_3 \frac{1}{3} e^{-(t_1 - t_m)}}{(1-\delta_2 - \delta_3)(1-e^{-(t_2-t_1)}) + (1-\delta_2 - \delta_3)\frac{1}{3}e^{-(t_2-t_1)} + \delta_2 \frac{1}{3}e^{-(t_2-t_m)} + \delta_3 \frac{1}{3}e^{-(t_1-t_m)}}$$

$f_{BC1_2}$

$$= \frac{\delta_2 \left(1 - e^{-(t_2-t_m)}\right)}{\delta_2(1-e^{-(t_2-t_m)}) + \delta_2 \frac{1}{3}e^{-(t_2-t_m)} + (1-\delta_2 - \delta_3)\frac{1}{3}e^{-(t_2-t_1)} + \delta_3(1-e^{-(t_1-t_m)}) + \delta_3 \frac{1}{3}e^{-(t_1-t_m)}}$$

$f_{BC2_2}$

$$= \frac{\delta_2 \frac{1}{3} e^{-(t_2-t_m)}}{\delta_2(1-e^{-(t_2-t_m)}) + \delta_2 \frac{1}{3}e^{-(t_2-t_m)} + (1-\delta_2 - \delta_3)\frac{1}{3}e^{-(t_2-t_1)} + \delta_3(1-e^{-(t_1-t_m)}) + \delta_3 \frac{1}{3}e^{-(t_1-t_m)}}$$

$f_{BC_1}$

$$= \frac{(1-\delta_2 - \delta_3)\frac{1}{3} e^{-(t_2-t_1)}}{\delta_2(1-e^{-(t_2-t_m)}) + \delta_2 \frac{1}{3}e^{-(t_2-t_m)} + (1-\delta_2 - \delta_3)\frac{1}{3}e^{-(t_2-t_1)} + \delta_3(1-e^{-(t_1-t_m)}) + \delta_3 \frac{1}{3}e^{-(t_1-t_m)}}$$

$f_{BC1_3}$

$$= \frac{\delta_3 \left(1 - e^{-(t_1-t_m)}\right)}{\delta_2(1-e^{-(t_2-t_m)}) + \delta_2 \frac{1}{3}e^{-(t_2-t_m)} + (1-\delta_2 - \delta_3)\frac{1}{3}e^{-(t_2-t_1)} + \delta_3(1-e^{-(t_1-t_m)}) + \delta_3 \frac{1}{3}e^{-(t_1-t_m)}}$$

$f_{BC2_3}$

$$= \frac{\delta_3 \left(1 - e^{-(t_1-t_m)}\right)}{\delta_2(1-e^{-(t_2-t_m)}) + \delta_2 \frac{1}{3}e^{-(t_2-t_m)} + (1-\delta_2 - \delta_3)\frac{1}{3}e^{-(t_2-t_1)} + \delta_3(1-e^{-(t_1-t_m)}) + \delta_3 \frac{1}{3}e^{-(t_1-t_m)}}$$

(Hibbins and Hahn 2019). To obtain the results in Figure 4 of the main text and Supplementary Figure 2, we calculated $h_{AB}|AB$ and $h_{BC}|BC$ over a wide range of parameter values, and asked which quantity was smaller in which parts of the parameter space.

**Supplementary Figures**

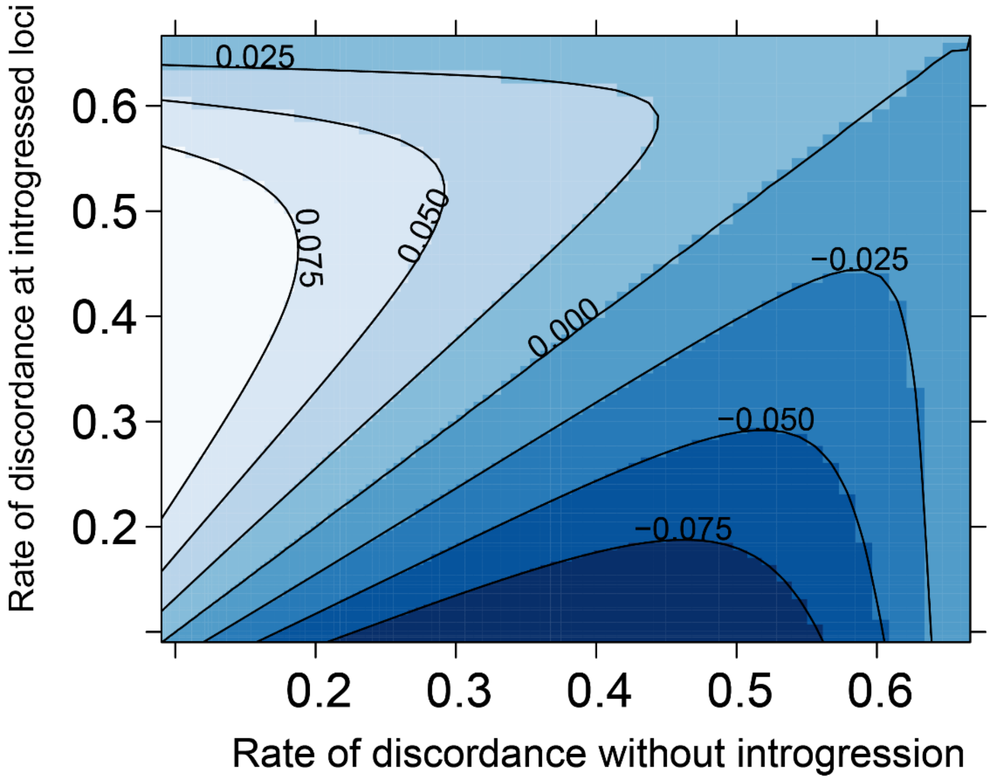


*Supplementary Figure 1*: Difference in the performance of quartet vs. parsimony methods for species tree inference in the presence of introgression. The contour values show the minimum amount of introgression necessary to mislead parsimony methods minus the minimum amount for quartet methods.

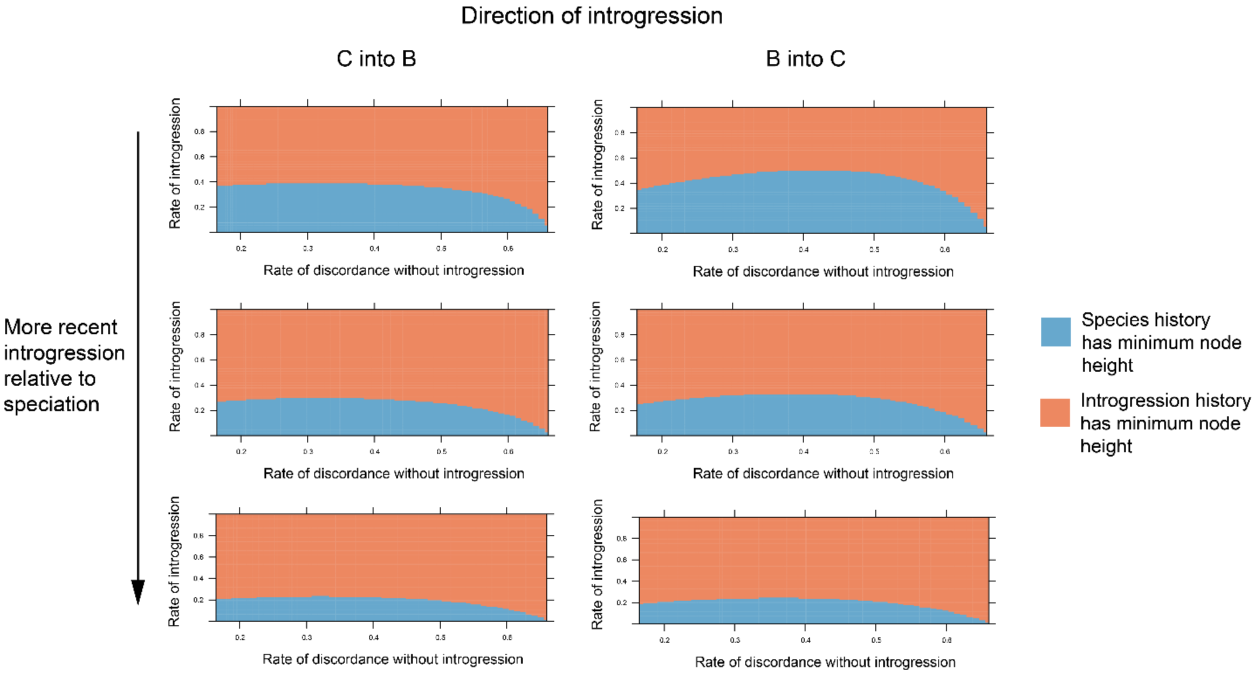


*Supplementary Figure 2*: Minimum node height over the parameter space of our model, with the two directions of introgression shown separately.

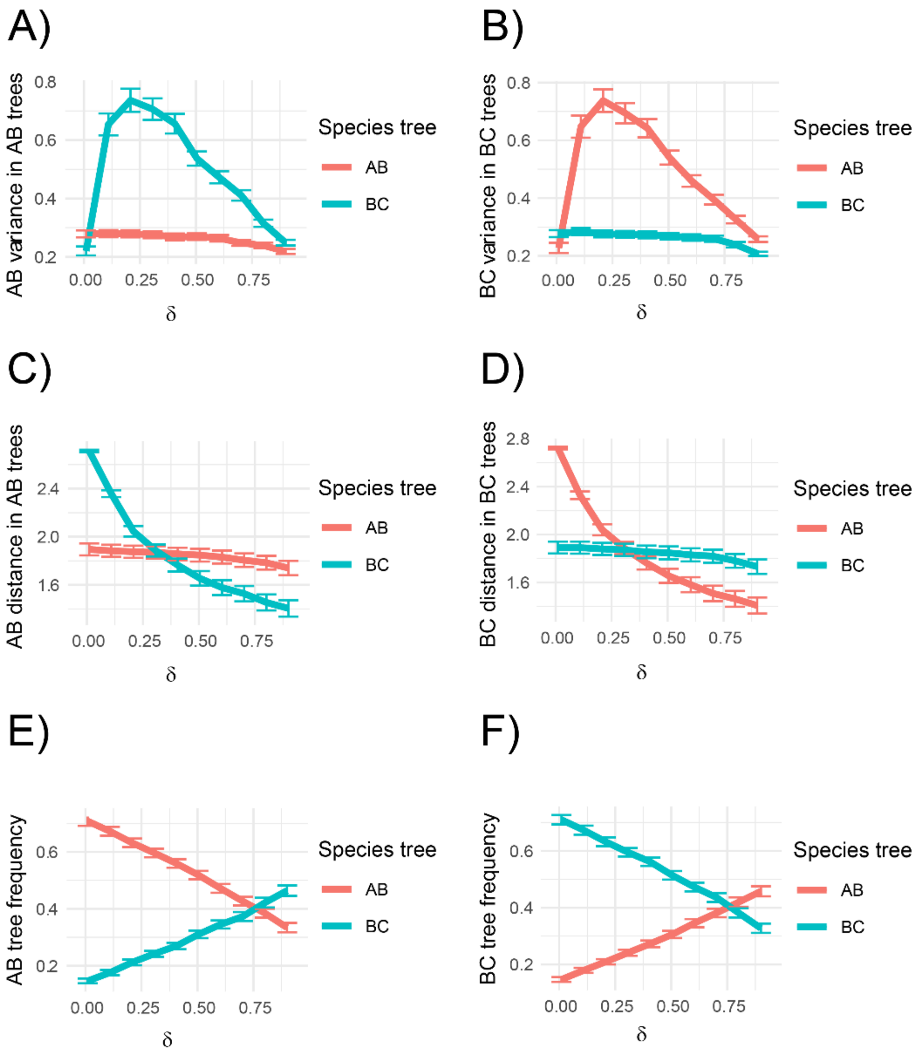


*Supplementary Figure 3*: Behavior of the six most informative features in our machine learning model, for P2 into P3 introgression (B into C or B into A, depending on which history is which).
